## Supplemental material for "Autoinhibition and regulation by phosphoinositides of ATP8B1, a human lipid flippase associated with intrahepatic cholestatic disorders"

### Supporting information

#### Figure supplements

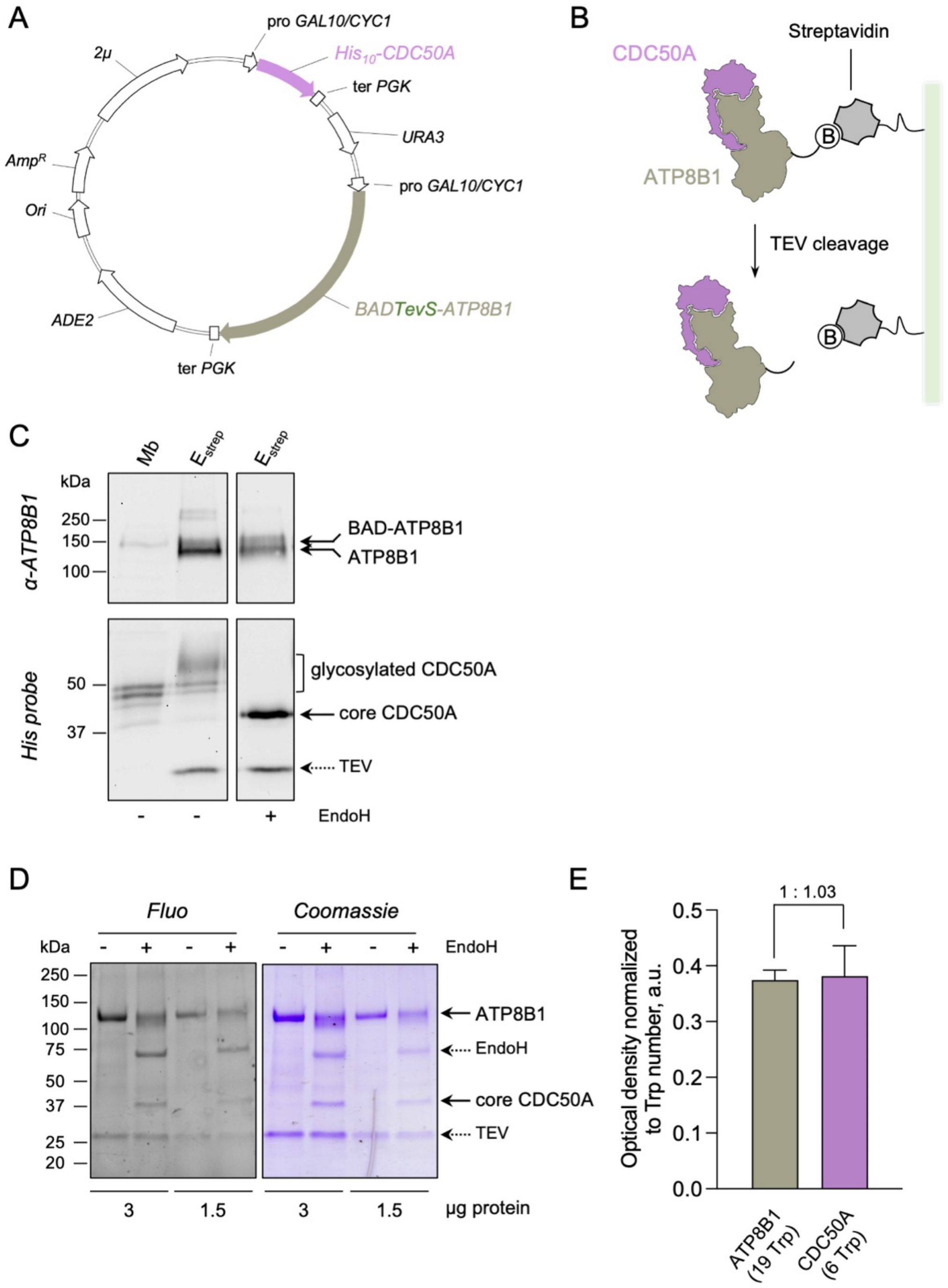


**Figure 1 – figure supplement 1: Strategy for purification of the ATP8B1-CDC50A complex.** *(A)* Map of the plasmid used for co-expression of BAD-ATP8B1 and $His_{10}$-CDC50A. Both ATP8B1 and CDC50A genes are cloned into the same expression vector. The cDNA sequence of human CDC50A is fused to a N-terminal deca-histidine tag ($His_{10}$), and the cDNA sequence of human ATP8B1 is fused to a N-terminal biotin acceptor domain (BAD) for *in vivo* biotinylation in yeast and further affinity purification on streptavidin beads (Jidenko et al., 2006). The BAD tag is followed by the sequence of a TEV protease cleavage site. ATP8B1 and CDC50A open reading frames are under the control of a strong hybrid galactose-inducible promoter (*GAL10-CYC1*). Ter PGK: sequence of the phosphoglycerate kinase used for termination of transcription; ADE2: auxotrophy selection marker for adenine; Ori: bacterial replication origin; AmpR: gene conferring resistance to ampicillin; 2µ: yeast replication origin; URA3: auxotrophic selection marker for uracil. *(B)* Purification scheme. The BAD-ATP8B1/$His_{10}$-CDC50A complex is solubilized from yeast membranes and applied onto streptavidin beads. Incubation with TEV protease allows elution of ATP8B1-CDC50A complex from the beads and removal of the BAD tag. *(C)* ATP8B1-CDC50A purification assessed by immunoblot analysis. Yeast crude membranes (Mb) and eluted proteins ($E_{strep}$) were detected with either an anti-ATP8B1 antibody (upper panel) or a histidine probe (lower panel). For deglycosylation experiments, the eluted fraction was treated with EndoH. The band for ATP8B1 appears rather diffuse upon EndoH treatment because boiling prior loading induced aggregation. *(D)* Purified ATP8B1-CDC50A complex was loaded at the indicated amount onto a haloalkane-containing 4-15% gradient SDS-PAGE for in-gel fluorescence analysis (left, Fluo) and subsequent Coomassie Blue staining (right, Coomassie). Assuming that all tryptophan residues react similarly to haloalkane in denaturing conditions, the fluorescence intensity measured accurately reflects the number of tryptophans in the protein and therefore the amount of protein loaded. *(E)* Relative fluorescence intensity of ATP8B1 and CDC50A bands observed in panel D, as deduced from quantification using ImageJ. The error bars represent the mean ± s.d. calculated from the loading of two different quantities of purified ATP8B1-CDC50A complex (3 and 1.5 $\mu$g). The experiment displayed in *(D)* and *(E)* is representative of three independent ones with similar results. Source files related to Figure 1 – figure supplement 1E are available in Figure 1 – figure supplement 1 – Source Data 1.

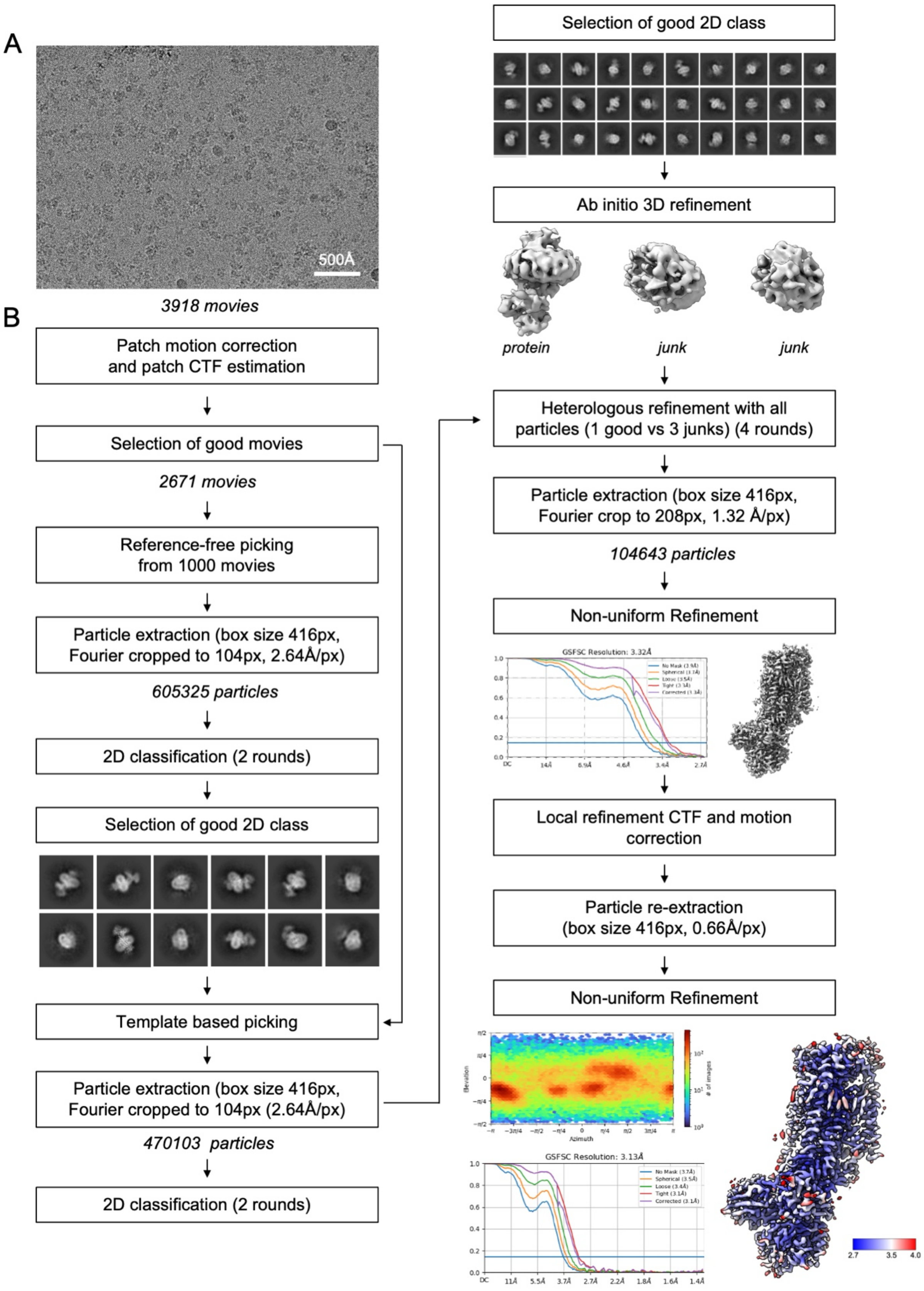


**Figure 2 – figure supplement 1: Data processing flow chart.**
*(A)* Representative motion-corrected and dose weighted micrograph. *(B)* Data-processing workflow performed in CryoSparc v3.

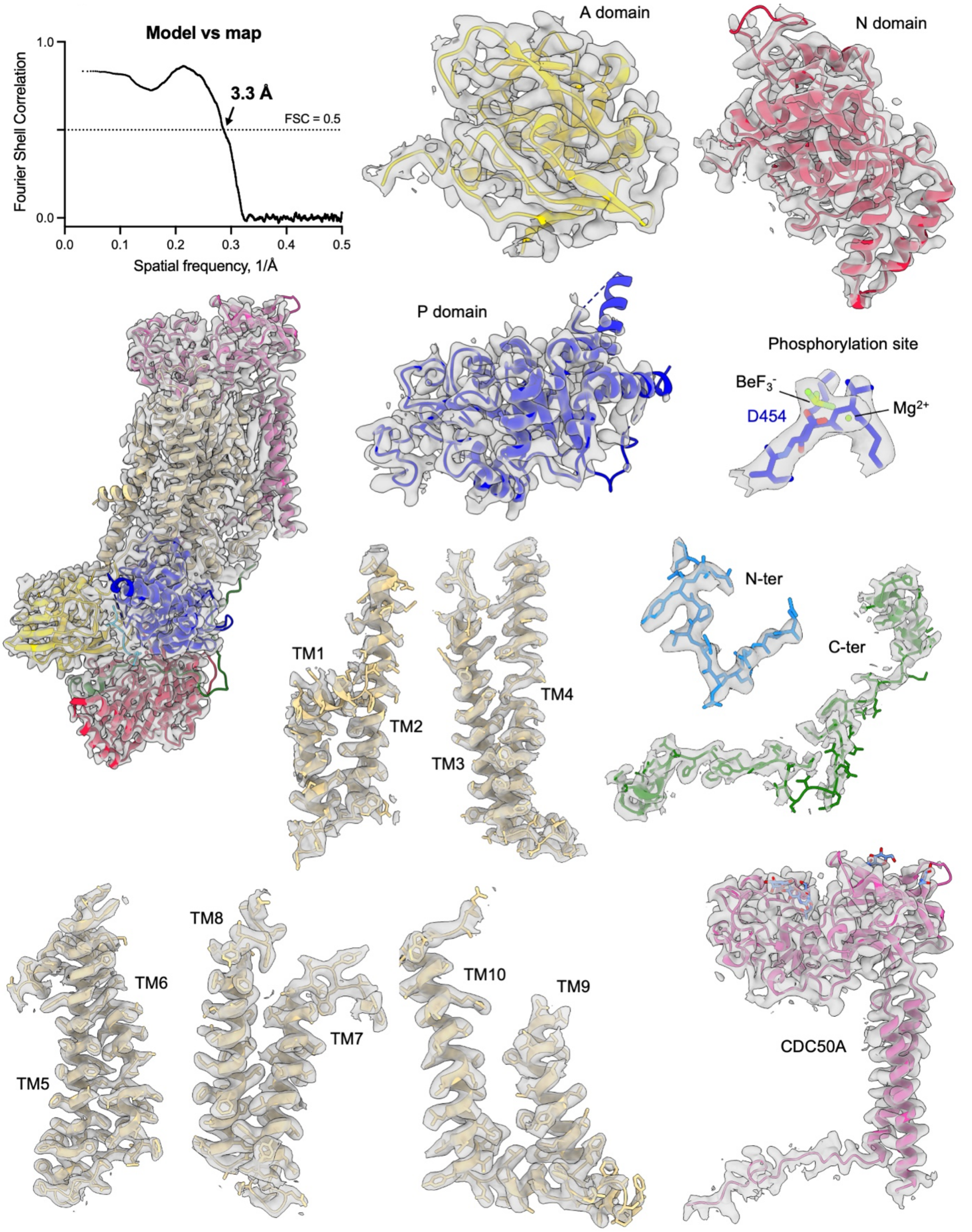


**Figure 2 – figure supplement 2: Cryo-EM density of the ATP8B1-CDC50A complex and its corresponding model.**
Map to model FSC curve and cryo-EM densities from different areas of the ATP8B1-CDC50A complex in the E2P autoinhibited state. Cryo-EM map contour levels used are 5 to 9. TM: transmembrane helical segments.

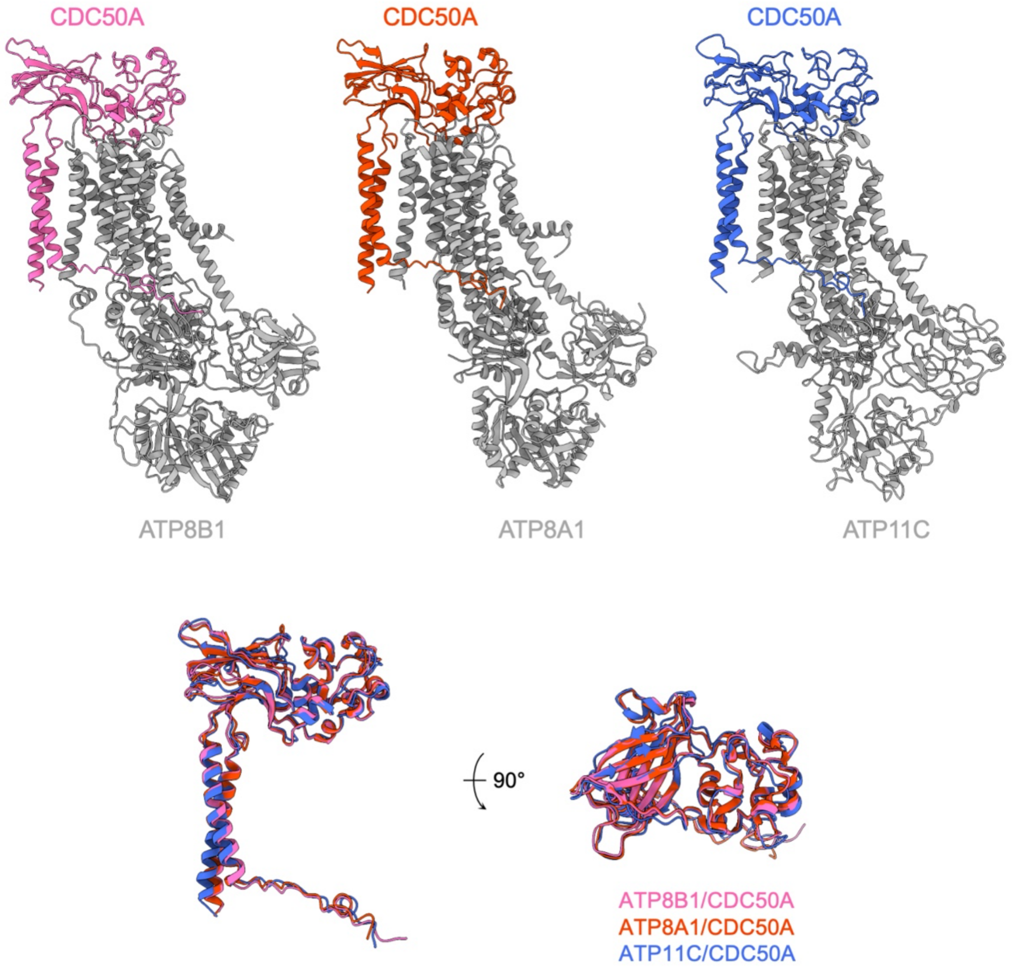


| CDC50A all-Cα RMSD (Å) | | | |
|---|---|---|---|
| | *ATP8B1/CDC50A* | *ATP8A1/CDC50A* | *ATP11C/CDC50A* |
| *ATP8B1/CDC50A* | - | *0.78* | *1.15* |
| *ATP8A1/CDC50A* | *0.78* | - | *1.14* |
| *ATP11C/CDC50A* | *1.15* | *1.14* | - |

**Figure 2 – figure supplement 3: Structural comparison of P4-ATPases-CDC50A complexes of known structure.**
*(A)* Atomic models of CDC50A in complex with ATP8B1, ATP8A1 (PDB: 6K7L) and ATP11C (6LKN), colored in pink, orange and blue respectively. P4-ATPases are colored in gray. *(B)* Structural alignment of CDC50A from 3 complexes formed with different human P4-ATPases and associated Cα-RMSD. Colors are as in *(A)*.

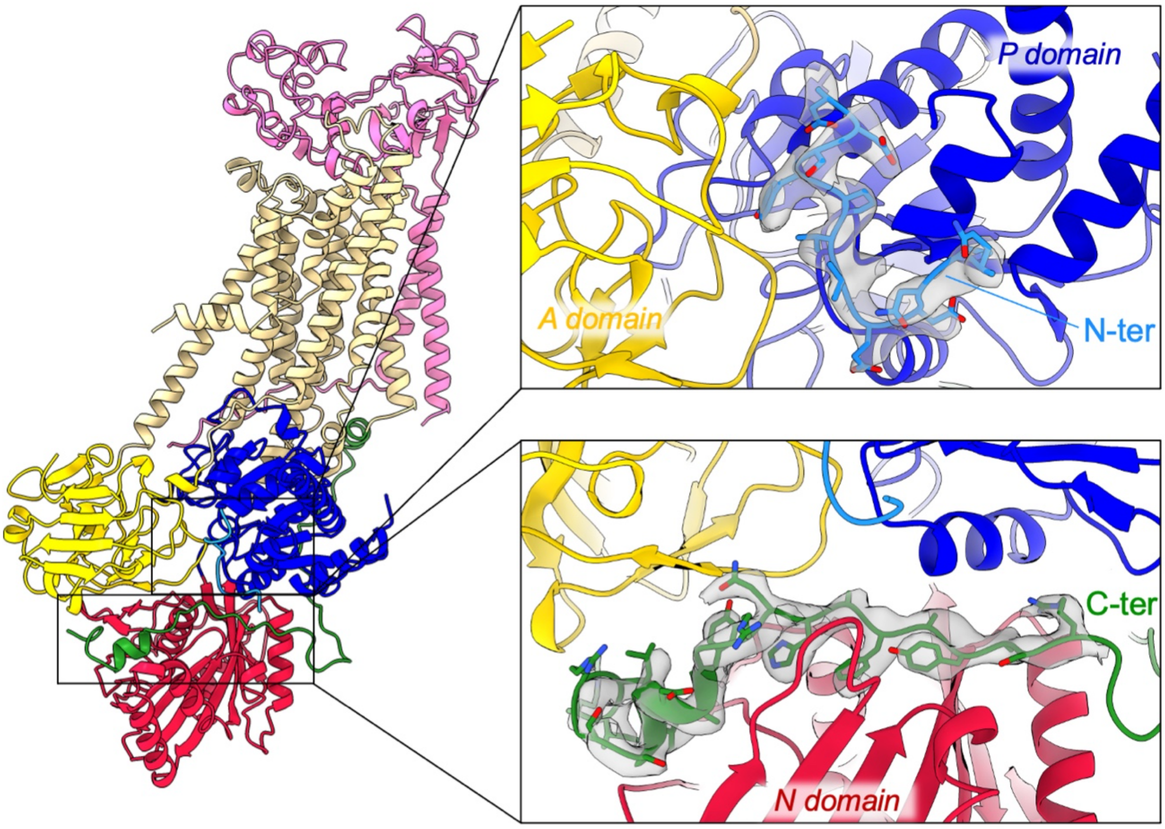


**Figure 3 – figure supplement 1: overall and close-up views of the N- and C-terminal extensions of ATP8B1 and their corresponding EM densities.**

The cytosolic A-, N- and P-domains of ATP8B1 are colored in yellow, red and blue, respectively. The transmembrane domain of ATP8B1 is colored in tan. The N- and C-terminal tails of ATP8B1 and their side chains are colored in cyan and green, respectively. CDC50A is colored in pink.

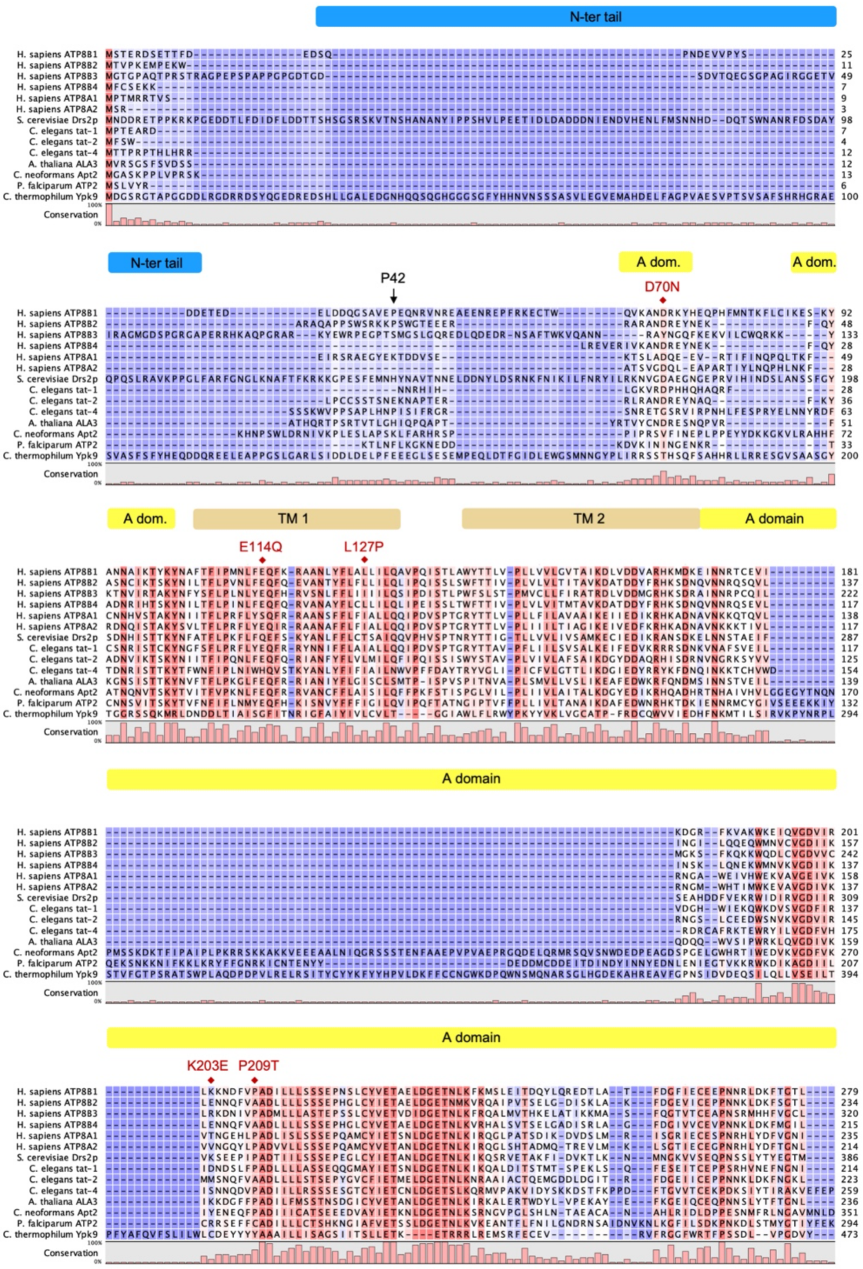


**Figure 4 – figure supplement 1: Sequence alignment of selected P4-ATPases**

Sequences of the human ATP8B1, ATP8B2, ATP8B3, ATP8B4, ATP8A1, ATP8A2, the *S. cerevisiae* Drs2, the *C. elegans* tat-1, tat-2, tat-4, the *A. thaliana* ALA3, the *C. neoformans* Apt2, the *P. falciparum* ATP2 were aligned based on the presence of the conserved motif found on the regulatory C-terminus of ATP8B1, Drs2, ATP8A1 and ATP8A2. For comparison, the sequence of the *C. thermophilum* P5-ATPase Ypk9 is shown. Sequences were aligned using the ClustalW server and manually edited for the C-terminal region because of the very low conservation of this region (apart from the (G/A)(Y/F)AFS motif). The shading indicates conservation (blue 0% – red 100%). Mutations found in PFIC1, BRIC1 and ICP1 are indicated with red diamonds. The various cytosolic domains and transmembrane helices of ATP8B1 are indicated above the sequences. Residues P42 and E1174, after which 3C protease cleavage sites were added, and the S1223 which is phosphorylated in mouse ATP8B1 are highlighted with arrows. The sequence used for the synthesis of the autoinhibitory C-terminal peptide is emphasized by a green box. Uniprot identifiers: ATP8B1 (O43520), ATP8B2 (P98198), ATP8B3 (O60423), ATP8B4 (Q8TF62), ATP8A1 (Q9Y2Q0), ATP8A2 (Q9NTI2),Drs2 (P39524), tat-1 (Q9U280), tat-2 (Q9TXV2), tat-4 (H2KZ37), ALA3 (Q9XIE6), Apt2 (Q5K6X2), PfATP2 (Q8I5L4), Ypk9 (G0S7G9).

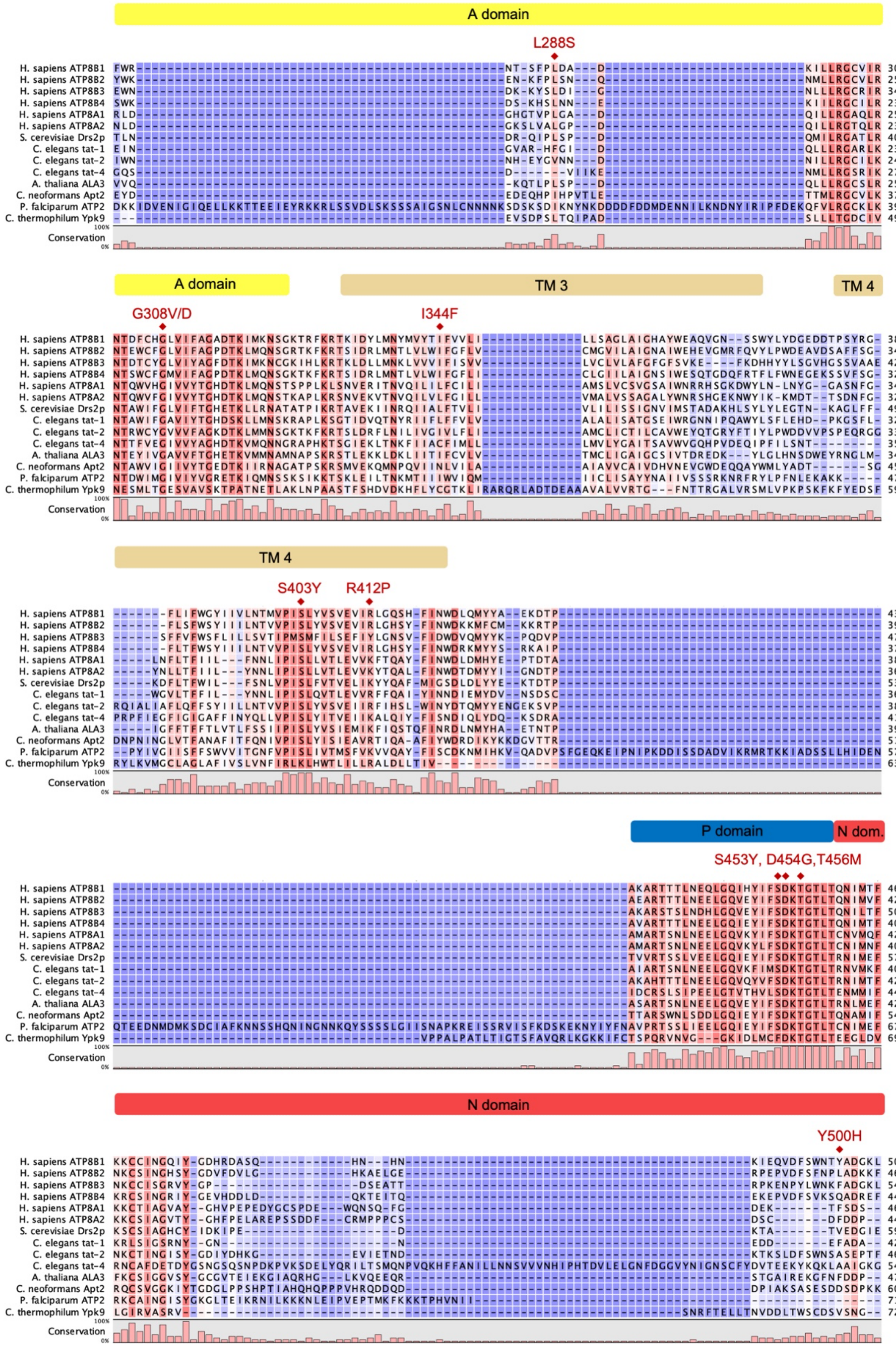


**Figure 4 – figure supplement 1: continued**

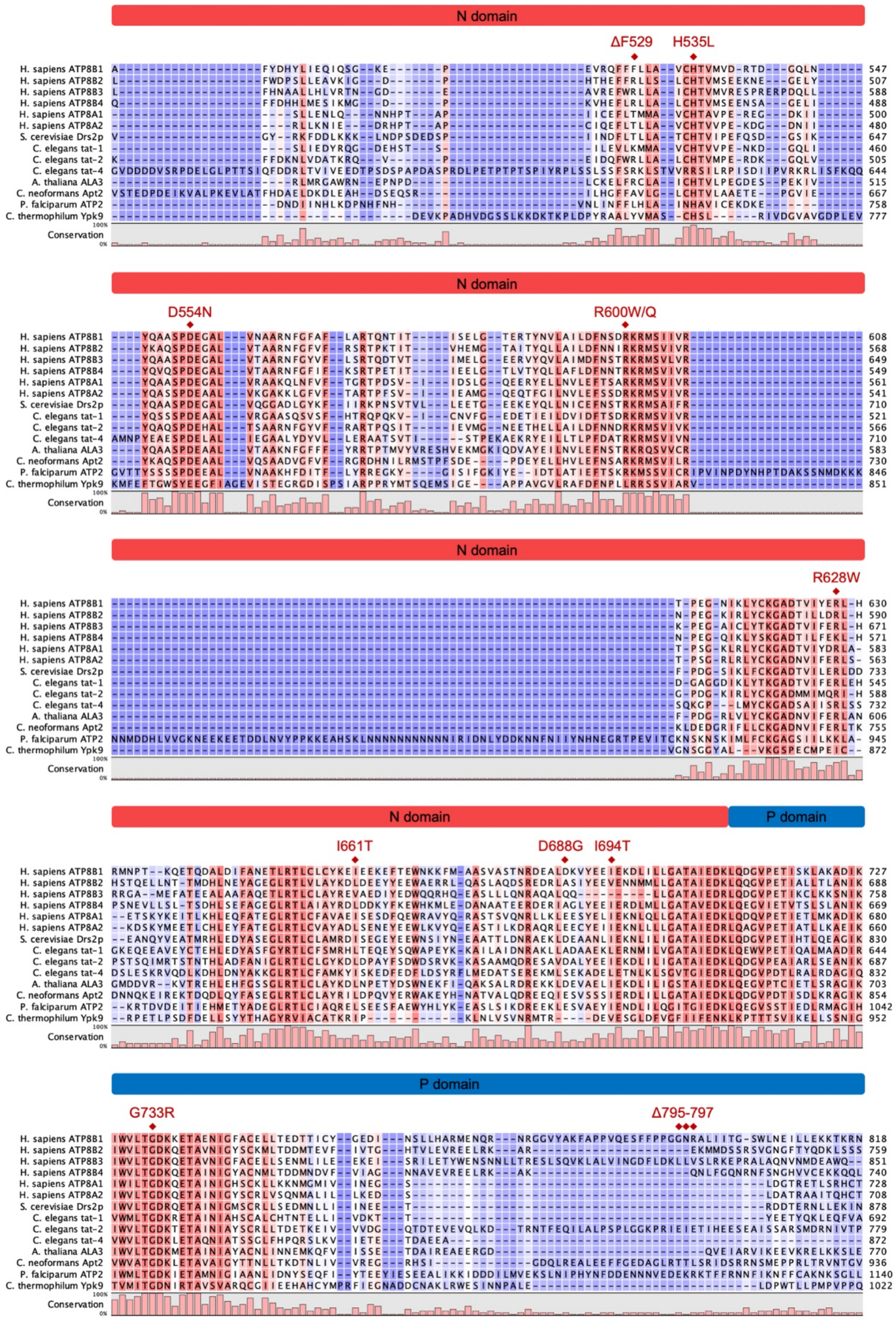


**Figure 4 – figure supplement 1: continued**

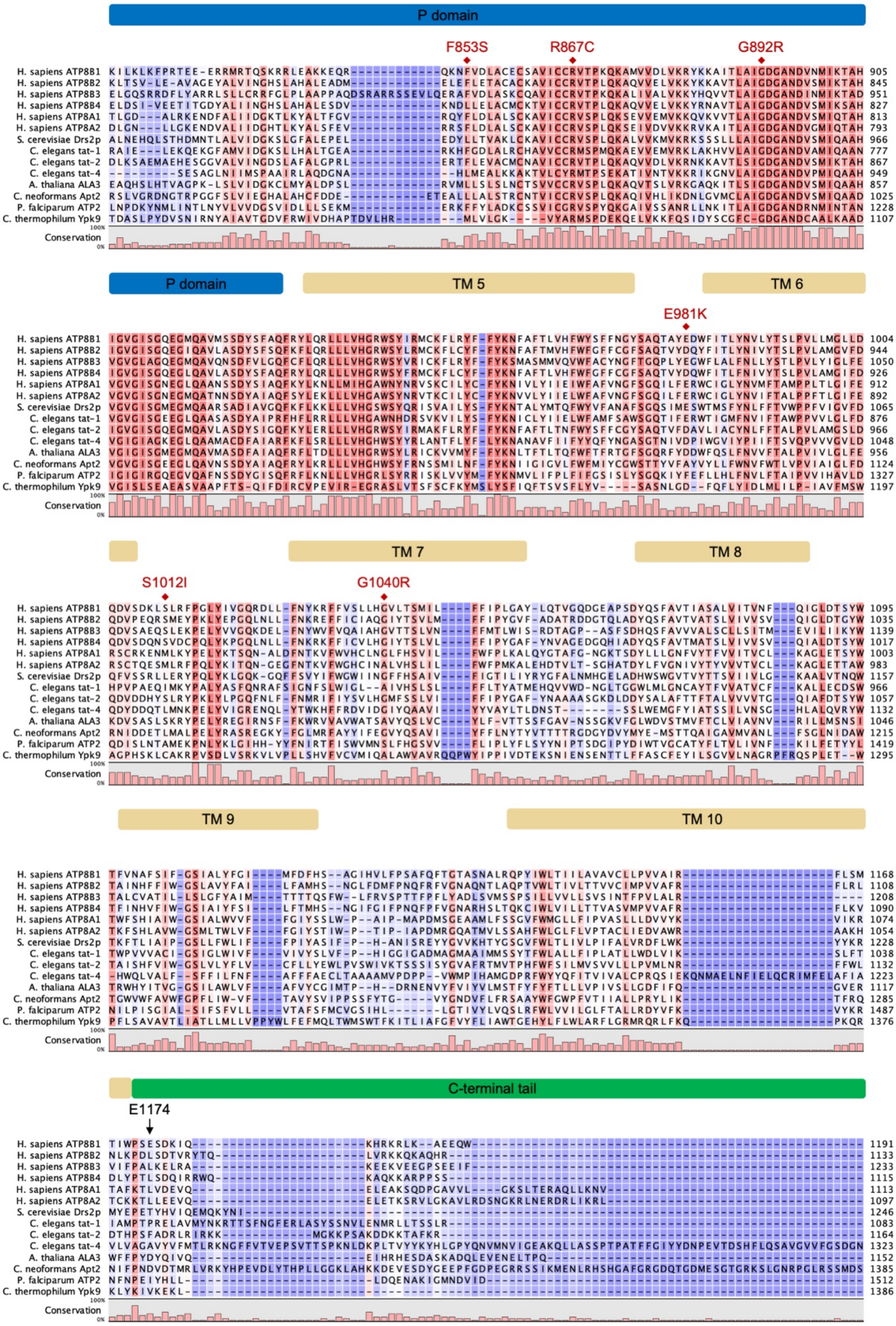


**Figure 4 – figure supplement 1: continued**

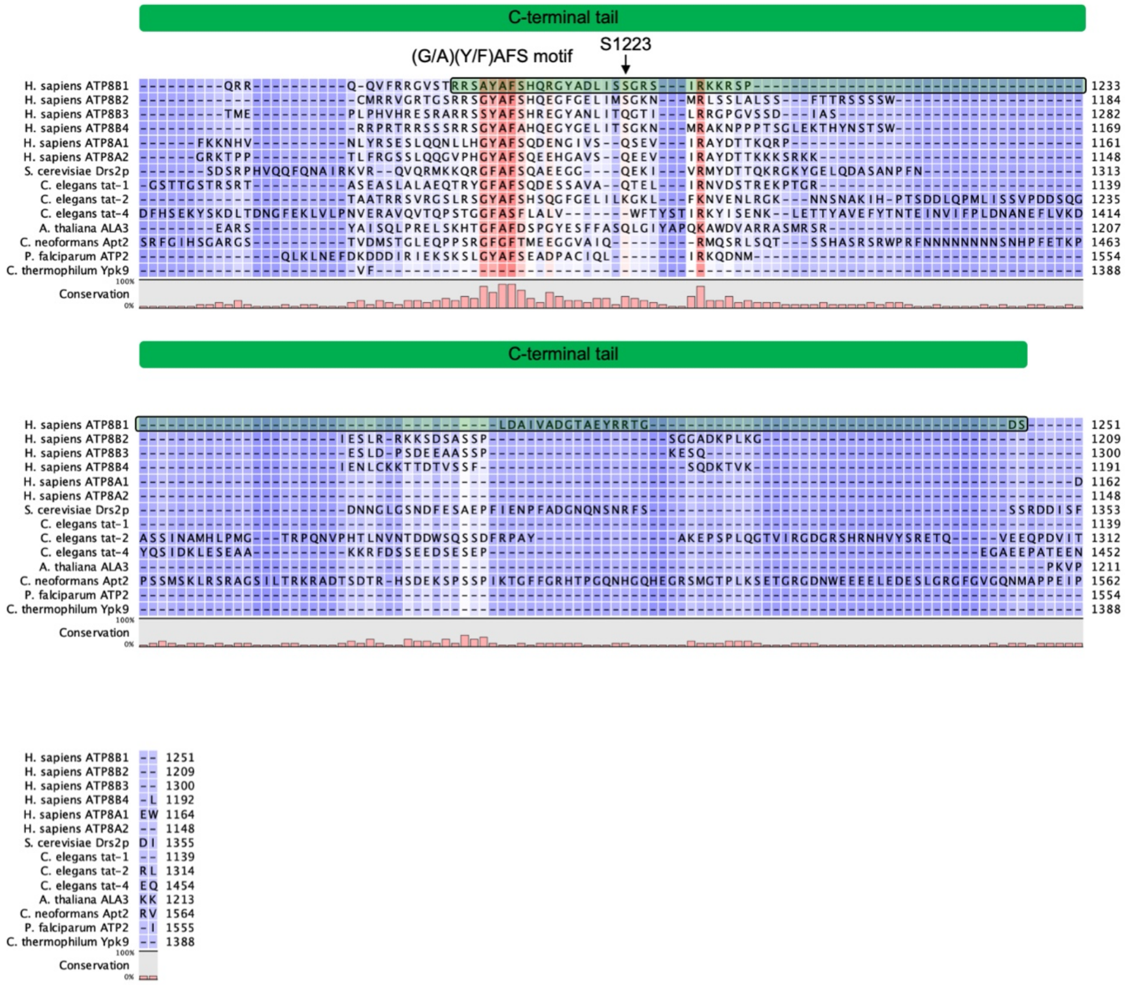


**Figure 4 – figure supplement 1: continued**

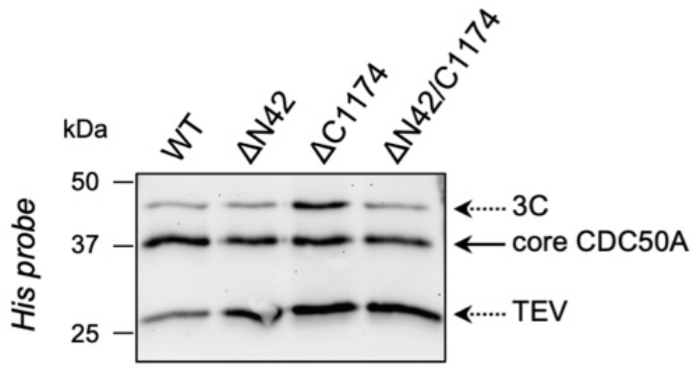


**Figure 4 – figure supplement 2: The amount of CDC50A which co-elutes with ATP8B1 upon on-column cleavage with TEV and 3C proteases is similar for wild-type ATP8B1 (WT) and the 3C protease cleavage site insertion mutants (ΔN42, ΔC1174, ΔN42/C1174**). The ATP8B1-CDC50A complex recovered from streptavidin beads upon proteolytic cleavage was denatured, treated with EndoH for 1 h at 37°C, and analyzed by immunoblotting with a Histidine probe.

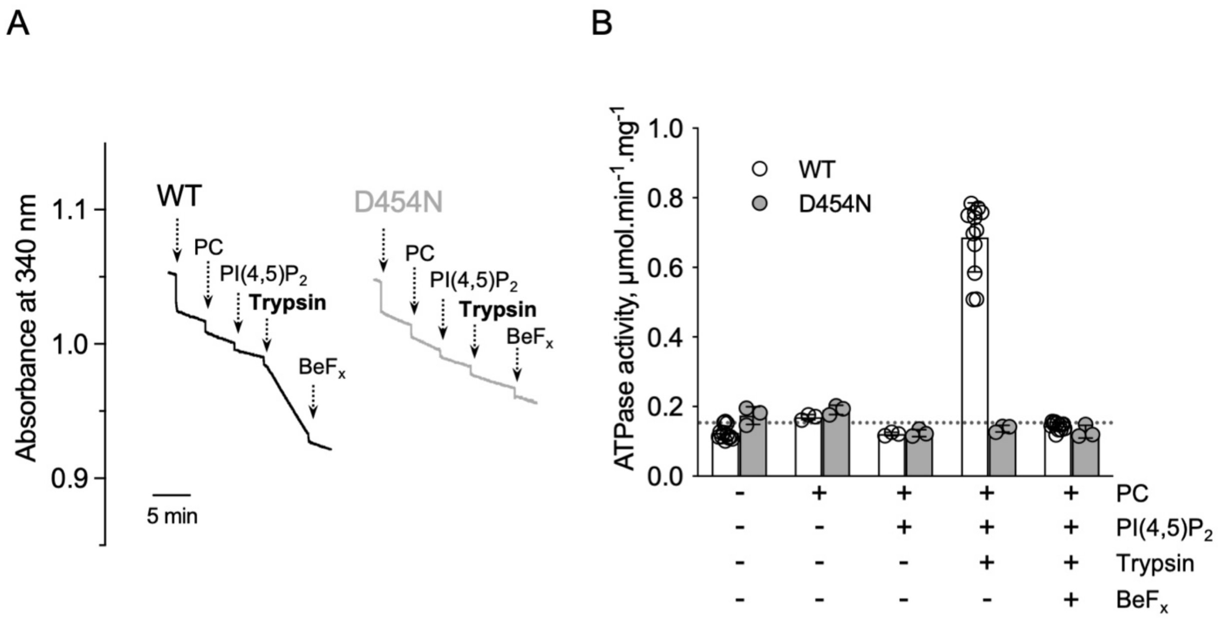


**Figure 4 – figure supplement 3: ATPase activity measurements of streptavidin-purified WT and catalytically-inactive D454N ATP8B1-CDC50A.** *(A)* ATPase activity of the purified ATP8B1-CDC50A complex determined in DDM/CHS at 30°C, using an enzyme-coupled assay, where the kinetics of NADH oxidation is monitored continuously. The various additions in the assay cuvette are indicated with arrows. ATP8B1 was added at ~ 2 µg ml$^{-1}$ to continuously stirred cuvettes in an assay medium containing 1 mM MgATP, 0.5 mg ml$^{-1}$ DDM, and 0.01 mg ml$^{-1}$ CHS in buffer B. PC and PI(4,5)$P_2$ were added at 0.1 mg ml$^{-1}$ and 0.025 mg ml$^{-1}$, respectively, resulting in a DDM final concentration of 1.25 mg ml$^{-1}$. Trypsin and $BeF_x$ were added at 0.07 mg ml$^{-1}$ and 1 mM, respectively. The rate of ATP hydrolysis corresponds to the slope measured after each addition. Activity is revealed upon addition of trypsin. *(B)* Specific ATPase activity of WT ATP8B1-CDC50A measured from traces such as that displayed in *(A)*. The dotted line represents the background NADH oxidation level, as measured before addition of ATP8B1-CDC50A in the assay cuvette. Data in (*B*) are a mean ± s.d. of 6 to 12 replicate experiments. PC: phosphatidylcholine. Source files related to Figure 4 – figure supplement 3B are available in Figure 4 – figure supplement 3 – Source Data 1.

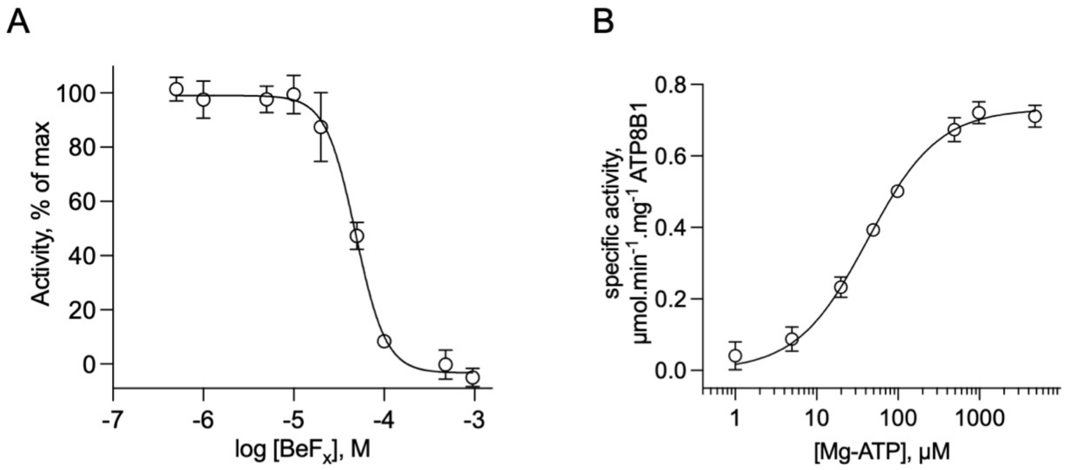


**Figure 4 – figure supplement 4: Catalytic properties of the purified ATP8B1-CDC50A complex.** *(A)* Sensitivity to beryllium fluoride of ATP8B1 turnover rate. ATPase activity of ΔN42/C1174 ATP8B1 was measured at 30°C in the presence of increasing concentrations of $BeF_x$, with 2 mM DDM, 115 $\mu$M PC, 23 $\mu$M PI(4,5)$P_2$, and ~0.5 $\mu$g $ml^{-1}$ ΔN42/C1174 in the assay cuvette. Data are a mean ± s.d. of 3 replicate experiments. The activity in the absence of $BeF_x$ was taken as 100% and data were fitted to an inhibitory dose-response equation with variable slope. 95% confidence interval: CI[$4.29x10^{-5}$, $5.43x10^{-5}$]. *(B)* Dependence on MgATP of the turnover rate. ATPase activity of ΔN42/C1174 ATP8B1 was measured at 30°C in the presence of increasing concentrations of MgATP, with 2 mM DDM, 115 $\mu$M PC, 23 $\mu$M PI(4,5)$P_2$, and ~0.5 $\mu$g $ml^{-1}$ ΔN42/C1174 in the assay cuvette. Data are a mean ± s.d. of 3 replicate experiments. The data were fitted to a Michaelis-Menten equation. For panels *(B)* and *(C)*, the rate of ATP hydrolysis was corrected for NADH photobleaching. Source files related to Figure 4 – figure supplement 4A and 4B are available in Figure 4 – figure supplement 4 – Source Data 1.

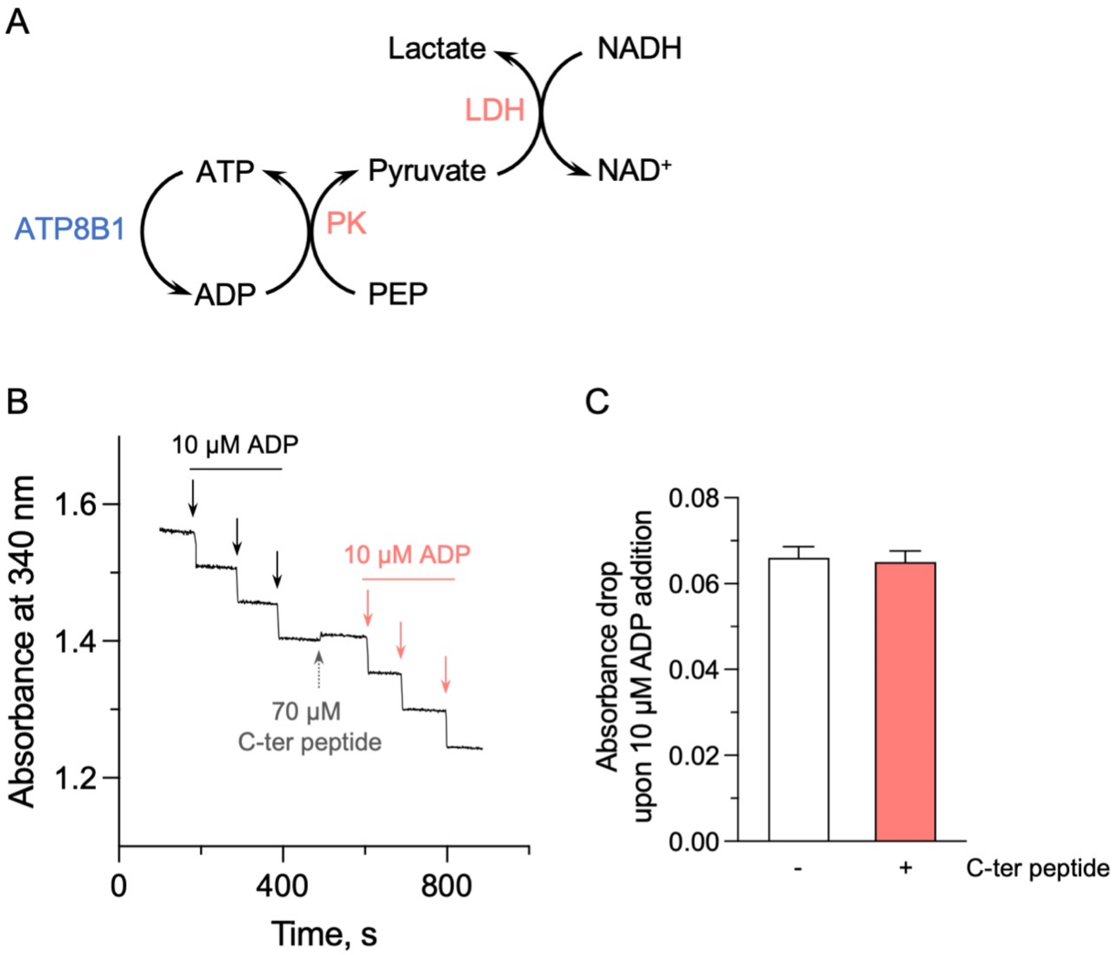


**Figure 5 – figure supplement 1: Effect of the ATP8B1 C-terminal peptide on the enzyme-coupled assay.**
*(A)* Schematic depicting the principle of the enzyme-coupled assay. PK: pyruvate kinase; LDH: lactate dehydrogenase. For each mole of ATP consumed by the ATP8B1-CDC50A complex, one mole of NADH is oxidized. This assay also allows continued regeneration of ATP from ADP. *(B)* Absorbance at 340 nm was monitored at 37°C, in the absence or presence of the ATP8B1 C-terminal peptide at 70 $\mu$M. Repeated additions of ADP at a final concentration of 10 $\mu$M (symbolized by each arrow) led to the expected oxidation of 10 $\mu$M NADH, resulting in a fast absorbance drop of ~0.06 AU at 340 nm. *(C)* Quantification of the absorbance decrease observed upon addition of 10 $\mu$M ADP, either in the absence (open bar) or in the presence (red bar) of the C-terminal peptide. Data are a mean ± s.d. of 3 technical replicates. Source files related to Figure 5 – figure supplement 1C are available in Figure 5 – figure supplement 1 – Source Data 1.

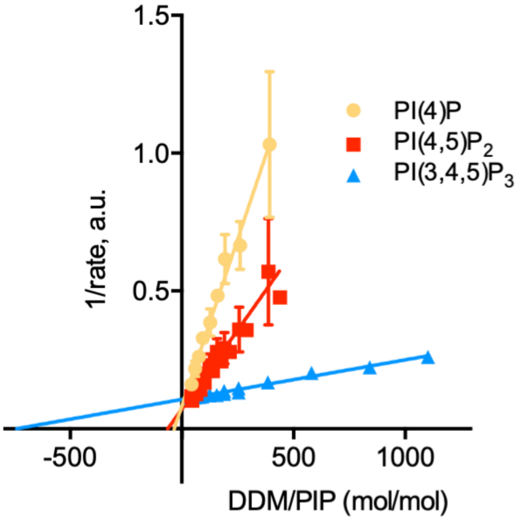


**Figure 6 – figure supplement 1: Determination of the kinetic parameters for activation of ATP8B1-CDC50A by PPIns.** Double reciprocal plot of data shown in Figure 6C. Source files related to Figure 6 – figure supplement 1 are available in Figure 6 – figure supplement 1 – Source Data 1.

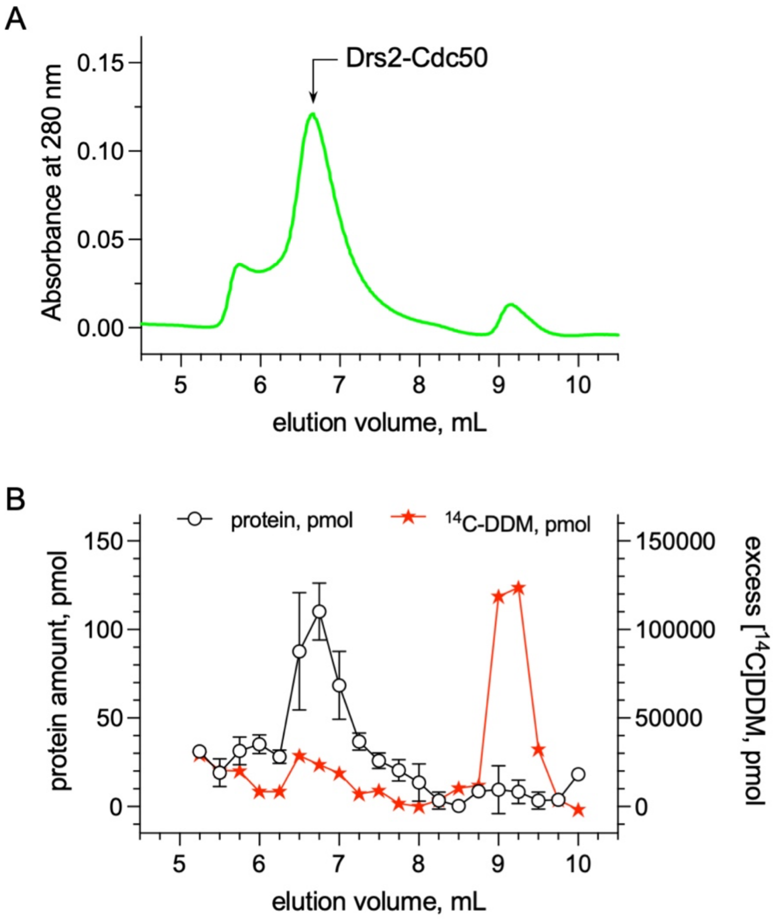


**Figure 6 – figure supplement 2: Quantification of the detergent bound to the transmembrane domain of Drs2-Cdc50.** *(A)* Size-exclusion chromatography (SEC) on a TSK3000 SW gel filtration column (Tosoh Bioscience) of DDM-purified Drs2-Cdc50 complex. The column was equilibrated with 50 mM MOPS-Tris pH 7, 100 mM KCl, 5 mM $MgCl_2$ supplemented with 0.5 mg mL$^{-1}$ DDM. The mobile phase also contained $^{14}$C-DDM to reach a specific activity of about 3.10$^{-5}$ $\mu$Ci per nmol of DDM. *(B)* Before loading affinity-purified Drs2-Cdc50 on the SEC column, at room temperature, the complex was incubated with $^{14}$C-DDM to reach a specific activity of 3.10$^{-5}$ $\mu$Ci per nmol of DDM. For each collected fraction, the protein and radioactive detergent content was quantified. The detergent/protein molar ratio of fractions collected for an elution volume of 6.5, 6.75 and 7 ml, where Drs2-Cdc50 peaks, was found to be around 272±56. Error bars represent the mean ± s.d. of 2 protein concentration measurements. Source files related to Figure 6 – figure supplement 2B are available in Figure 6 – figure supplement 2 – Source Data 1.

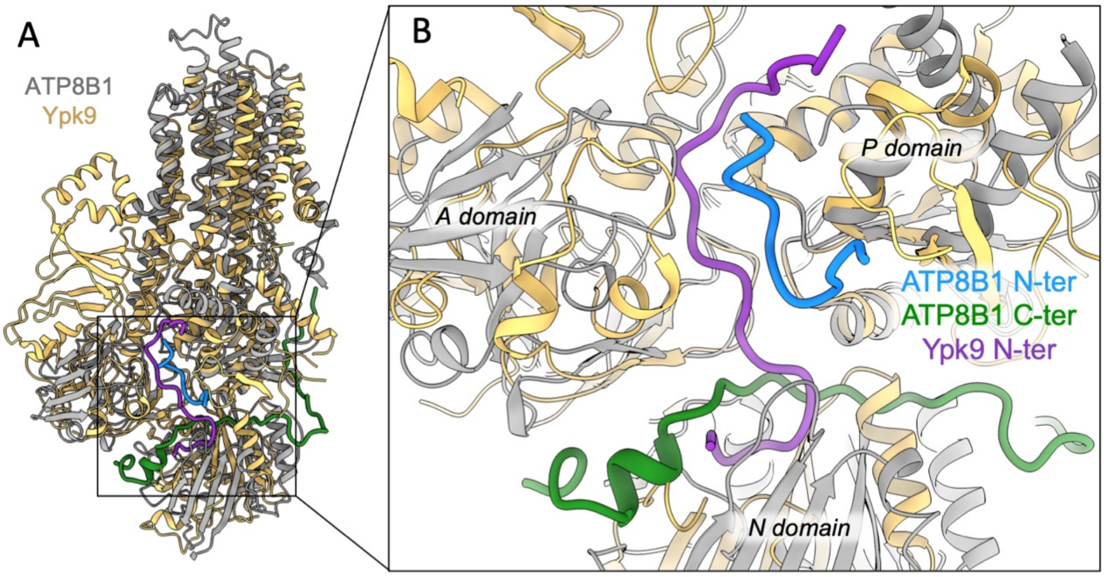


**Figure 7 – figure supplement 1: Structural comparison of ATP8B1 and Ypk9 autoinhibition mechanism.**

*(A)* Structural alignment of Ypk9 (PDB: 7OP8; yellow) and ATP8B1 (gray), both in the E2P inhibited state. *(B)* Close-up view of the region where the Ypk9 N-terminal tail (purple) and ATP8B1 N- and C-terminal tails (cyan and green, respectively) bind.
